# Mumemto: efficient maximal matching across pangenomes

**DOI:** 10.1101/2025.01.05.631388

**Authors:** Vikram S. Shivakumar, Ben Langmead

## Abstract

Aligning genomes into common coordinates is central to pangenome analysis and construction, but it is also computationally expensive. Multi-sequence maximal unique matches (multi-MUMs) are guideposts for core genome alignments, helping to frame and solve the multiple alignment problem. We introduce Mumemto, a tool that computes multi-MUMs and other match types across large pangenomes. Mumemto allows for visualization of synteny, reveals aberrant assemblies and scaffolds, and highlights pangenome conservation and structural variation. Mumemto computes multi-MUMs across 320 human genome assemblies (960GB) in 25.7 hours with under 800 GB of memory, and over hundreds of fungal genome assemblies in minutes. Mumemto is implemented in C++ and Python and available open-source at https://github.com/vikshiv/mumemto.

## 1 Introduction

Recent pangenomes can span hundreds of genomes. The Human Pangenome Reference Consortium, for example, has released hundreds of high-quality human genome assemblies [1]. Large pangenomes can shed light on sequence conservation and large-scale variants, and they provide reference panels for read alignment, genotype imputation, and sequence classification. As a result, there is a growing need for algorithms to align large pangenomes and reveal their underlying coordinate systems. This has spurred a range of new alignment and classification tools, as well as a new set of compressed-space algorithms for efficient construction of pangenome indexes.

Maximal exact matches (MEMs) and maximal unique matches (MUMs) are used in whole genome alignment [2], [3] and multiple sequence alignment [4]–[8] as syntenic anchors between more variable sequences. Collinear multi-MUMs can span large sections of conserved sequence in an MSA. However, existing methods for computing multi-MUMs do not scale well beyond relatively small collections of bacterial genomes. While tools like MUMmer4 [2] compute pairwise MUMs, the problem of aligning pangenomes is inherently multi-way, and computing them by way of pairwise alignments can require a quadratic number of sequence comparisons, plus a substantial merging step.

We introduce Mumemto, a tool to compute maximal exact or unique matches across many sequences. Mumemto uses prefix-free parsing (PFP) [9], a compressed-space method for computing the enhanced suffix array in sublinear space for pangenome sequence collections. For instance, Mumemto can compute multi-MUMs across 89 human genomes in under 4 hours, and across hundreds of fungal genomes in 2 minutes. We show that multi-MUMs can form a rudimentary MSA, reveal genomic conservation and structure, and identify aberrant assembly artifacts and potential pangenome issues. We propose Mumemto as a first-step pangenome diagnostic and visualization tool that scales efficiently for future large pangenome collections.

## 2 Results

### 2.1 Overview

Mumemto computes multi-MUMs and MEMs in a streaming algorithm over the suffix array (SA), Burrows-Wheeler Transform (BWT), and longest common prefix (LCP) arrays (Algorithm 1). Prefix-free parsing [9], computes these arrays efficiently for large, repetitive sequence collections like pangenomes. PFP computes these arrays sequentially, allowing Mumemto to consume them in a streaming fashion and avoiding the need for multiple passes or writing the arrays to disk. In short, Mumemto finds all relevant matches by performing some modest additional computation on top of the indexing process already used to produce compressed indexes like the *r*-index [10], [11] and move structure [12], [13].

Mumemto computes a variety of match types. Multi-MUMs are maximal exact matches that occur exactly once in all genomes. The other match types relax these constraints in some way; e.g. a partial multi-MUM may occur only some sequences (Figure 1).

**Figure 1:**
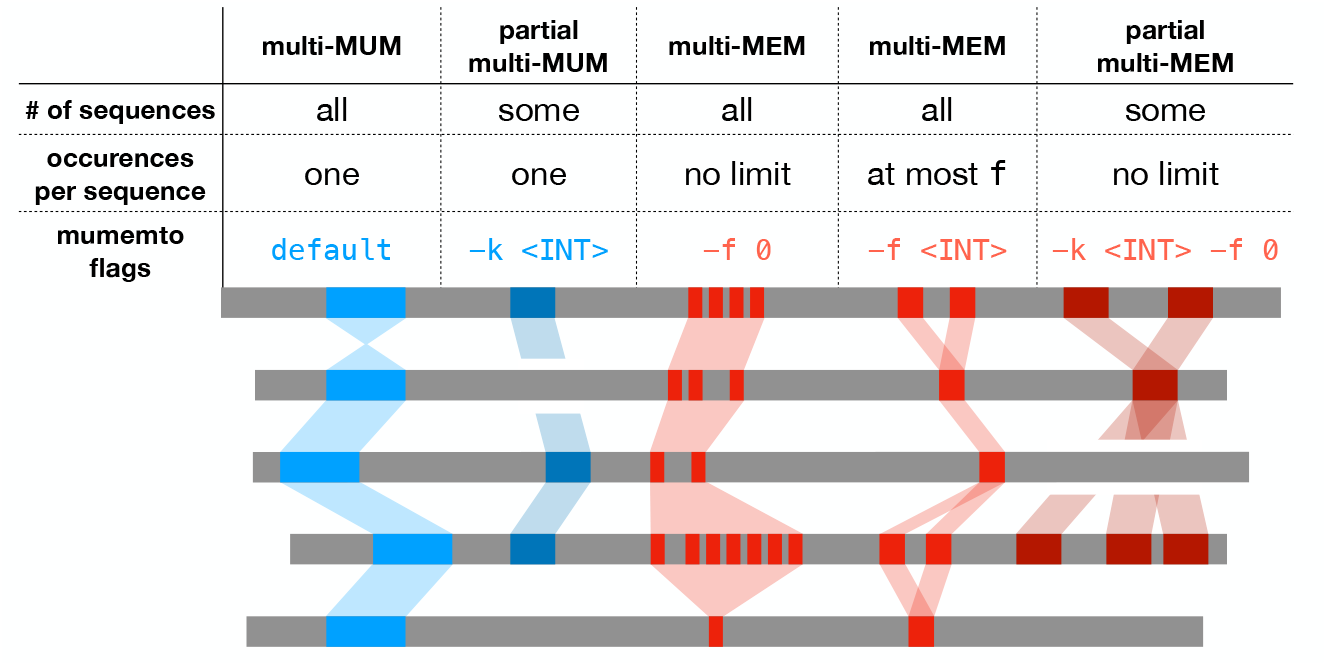
Exact match types that Mumemto can compute. Two flags to control how many sequences a match appears in (`-k`) and how many times a match may appear in any given sequence (`-f`).

Matches can be combined to define longer blocks of synteny, which may straddle important variants such as structural variants (SVs), e.g. inversion polymorphisms and rearrangements. The output of Mumemto can be used to visualize synteny and SVs by way of multi-MUMs. It can also characterize highly repetitive regions using multi-MEMs (Figure 5). As described below, Mumemto can also use partial-MUMs to identify potential assembly errors and other large-scale aberrations.

Mumemto differs from existing MUM-finding methods such as MUMmer4 [2] in that it computes matches shared across many sequences rather than just pairs of sequences. Though MUMs across *N* sequences could be computed by invoking a pairwise matching algorithm *O*(*N*^2^) times and merging the resulting pairwise MUMs, the quadratic time requirement is impractical for large pangenomes. For example, running MUMmer4 on all pairs of chr19 haplotypes from HPRC takes *>*30 hrs (30X slower than Mumemto), not counting the time to merge pairwise MUMs into multi-MUMs. As a result, we consider multi-MUM finding to be a distinct problem and omit pairwise methods from further comparisons.

### 2.2 Efficient core genome alignment and pangenome construction Mumemto is the fastest multi-MUM finder

To evaluate its multi-MUM finding algorithm, we compared Mumemto to the widely-used multi-MUM based multiple sequence aligners, Parsnp2 [5] and ProgressiveMauve [15]. Both tools find multi-MUMs as an intermediate step prior to collinear blocking and detailed alignment. Mauve uses a hash table of short seed matches, and extends unique seeds present in all sequences. Parsnp builds a compressed suffix graph over a reference sequence, then computes and merges multi-MUMs in each sequence across the collection. We found that both tools tend to miss a few multi-MUMs and falsely report a small number of non-unique matches, however these differences were negligible for the purposes of method evaluation.

We computed multi-MUMs across 89 haplotypes of each autosomal chromosome from the Human Pangenome Reference Consortium [1]. We measured the time and memory usage for only the MUM-finding step of each tool in Figure 2A-B. Mumemto was 7-11X faster than Mauve and 3-15X faster than Parsnp, while using 24%-44% less memory than Mauve and 39%-52% less than Parsnp. (Note that Mumemto was run on a single thread, while Mauve and Parsnp were run on 48 threads.) One exception was memory usage for chromosome 9, where Mumemto’s prefix-free parse included (by chance) long, unique centromeric substrings which are difficult to compress, yielding a slightly higher memory footprint compared to Parsnp and Mauve for that chromosome.

**Figure 2:**
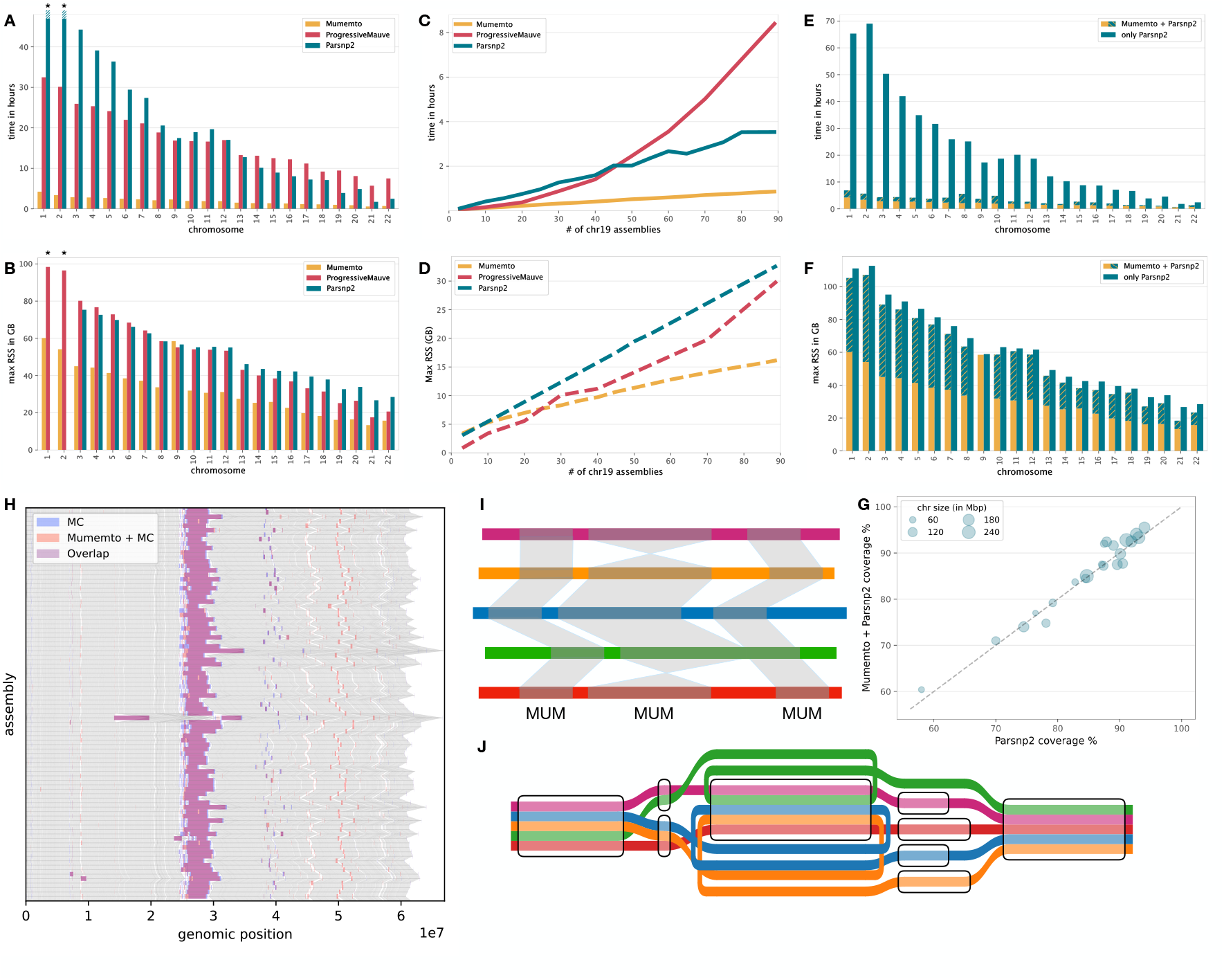
**(A-B)** Comparison of runtime and peak memory usage between multi-MUM finders. Only the initial multi-MUM computation was considered for ProgressiveMauve and Parsnp2. Note: Parsnp2 took >48 hrs for chromosome 1 and 2, so these are omitted. Parsnp2 and ProgressiveMauve were run with 48 threads, while Mumemto was run single-threaded. **(C-D)** Time and memory scaling comparison for increasing sequence collection sizes of chr19. **(E-F)** Comparison of time and memory for a Mumemto-seeded Parsnp2 alignment pipeline compared to the original Parsnp2 pipeline, and **(G)** a comparison of the alignments from each pipeline. **(H)** Regions excluded from Minigraph-Cactus (MC) while aligning chr19 assemblies, compared to regions excluded by a Mumemto-seeded MC pipeline (overlaid on a MUM synteny plot in gray). **(I-J)** Syntenic view of MUMs vs tube map [14] view of the equivalent graph.

When computing multi-MUMs over increasingly large collections of chromosome 19 haplotypes, Mumemto’s speed and memory scaled better than that of Mauve or Parsnp (Figure 2C-D), owing to the PFP algorithm’s ability to scale with the amount of non-redundant sequence. Further, we computed multi-MUMs across 320 human genome assemblies from HPRC (available at [16]) using 8 threads, which completed in 25.7 hours while using 800 GB of memory. If run serially, Mumemto would compute these multi-MUMs in under a week within 139 GB of memory.

#### Mumemto accelerates core genome alignment

Parsnp [17] uses multi-MUMs as initial guideposts to build a “core genome alignment,” i.e. a multiple alignment involving the conserved portions of the genomes. We modified the Parsnp pipeline to use Mumemto-computed multi-MUMs. We compared the original pipeline with the Mumemto-accelerated pipeline (Figure 2E-G). Though the peak memory footprint is dominated by the downstream portions of the Parsnp pipeline, the total runtime is up to 12× faster using Mumemto, while the overall alignment coverage is nearly identical to that of the original pipeline (Fig. 2G).

#### multi-MUMs form preliminary graphs

Multi-MUMs can also inform the construction of a preliminary pangenome graph. Collinear multi-MUMs represent conserved stretches of columns in the underlying MSA, and so are prime candidates for being collapsed into pangenome graph nodes. Gaps between collinear MUMs due to genomic variation are often short (<5bp), and common across haplotypes. For example, we found that among haplotypes of chr19, 78% of gaps between collinear MUMs were single nucleotide variants. As a result, for intraspecific pangenomes (such as HPRC) with high genome similarity, Mumemto can simplify and accelerate graph construction.

As a proof of concept, we compared various graph building strategies over HPRC haplotypes of chromosome 19. We built graphs using Mumemto and its reported multi-MUMs, and compared these to a graph built entirely with Minigraph-Cactus [18] (Table 1). The `Mumemto-full` strategy first computes multi-MUMs, then “collapses” gaps between collinear and adjacent multi-MUMs if the gap sequence is identical between any haplotypes. This strategy only includes small (≤ 100kb) gaps, assuming that larger gaps are similar to the “brnn” regions (e.g. centromeres), which are also excluded from the HPRC pangenome. The regions excluded in Minigraph-Cactus and `Mumemto-full` are further compared in Figure 2H. Mumemto does not perform base-level alignment in order to resolve variable-length gaps, but still achieves a compression ratio of 13.7X, meaning the 89-haplotype pangenome collapses to about 6.5 haplotypes worth of sequence needed to label the pangenome graph. As seen in Table 1, the `Mumemto-full` graph is larger than the others in terms of total sequence labels, its number of nodes and edges, and its memory footprint.

**Table 1:**
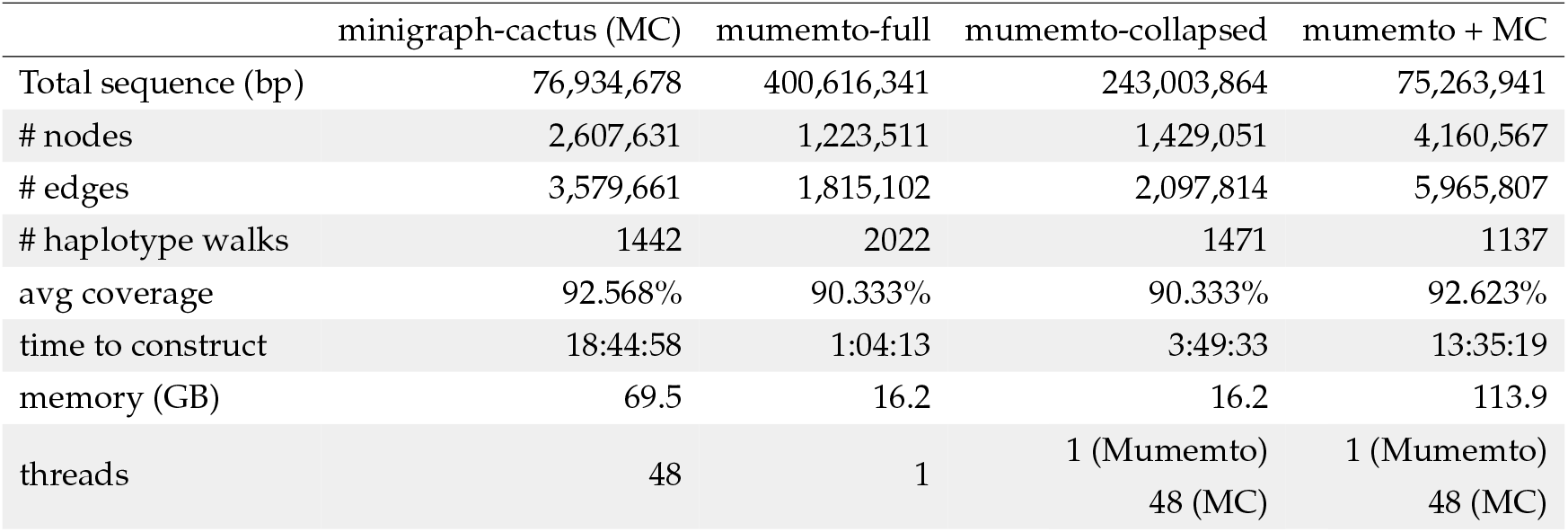
Comparison of different graph construction methods. `Mumemto-full` refers to a purely MUM-based graph where identical gaps between collinear MUMs are merged when possible. `Mumemto-collapsed` refers to a further compressed version of `Mumemto-full`, where large gaps between collinear MUMs are merged using Minigraph-cactus. `Mumemto + MC` refers to a Minigraph-cactus pipeline where the initial SV-only graph construction step is replaced with a simplified version of `Mumemto-full`.

|  | minigraph-cactus (MC) | mumemto-full | mumemto-collapsed | mumemto + MC |
| --- | --- | --- | --- | --- |
| Total sequence (bp) | 76,934,678 | 400,616,341 | 243,003,864 | 75,263,941 |
| # nodes | 2,607,631 | 1,223,511 | 1,429,051 | 4,160,567 |
| # edges | 3,579,661 | 1,815,102 | 2,097,814 | 5,965,807 |
| # haplotype walks | 1442 | 2022 | 1471 | 1137 |
| avg coverage | 92.568% | 90.333% | 90.333% | 92.623% |
| time to construct | 18:44:58 | 1:04:13 | 3:49:33 | 13:35:19 |
| memory (GB) | 69.5 | 16.2 | 16.2 | 113.9 |
| threads | 48 | 1 | 1 (Mumemto)<br>48 (MC) | 1 (Mumemto)<br>48 (MC) |

The `Mumemto-full` graph is larger because no attempt is made to collapse the interstitial sequence in large (>10kb) gaps between collinear MUMs. To address this, we identified the 50 largest gaps and aligned the inter-MUM sequence with Minigraph-Cactus. This graph (`Mumemto-collapsed`) is just under half the size of `Mumemto-full`, while only requiring an additional 2.75 hrs to compute.

We also considered how Mumemto could accelerate the Minigraph-Cactus pipeline. We replaced the initial SV graph construction step of Minigraph-Cactus with a simplified version of the `Mumemto-full` graph that included only short gaps (50bp-100kbp) that were partially shared (*<*45 unique gap sequences). We seeded the Minigraph-Cactus pipeline with this MUM-based SV graph, resulting in the `Mumemto + MC` graph. This strategy was faster than running Minigraph-Cactus, and provided a graph with a comparable coverage and compression ratio to Minigraph-Cactus (Table 1).

Finally, we compared the computational efficiency and accuracy of short read alignment to each of these graphs using giraffe [19] and Illumina reads from the HG002 individual from the Google Brain dataset [20]. We found comparable alignment quality and speed (Table S1). However, we noted that the Mumemto-seeded MC graph was slower for alignment, likely due to its larger graph. Nonetheless, the fast construction time and comparable sequence compression represents the potential for a Mumemto-accelerated approach for constructing pangenome graph indexes.

### 2.3 Mumemto reveals aberrations in pangenome assemblies

We found that examining the collinear MUMs reported by Mumemto revealed and helped to visualize aberrant features of pangenome assemblies. Large, private insertions and deletions in a single sequence manifested as a characteristic pattern of short, spurious MUMs spread across the genome (Figure 3A-B). These spurious MUMs are collinear in all but one sequence, helping to pinpoint the affected region (Figure 3D-E). If a large number of collinear MUM pairs are separated in a specific sequence, Mumemto can identify the problem region as either an insertion (Figure 3A) or deletion (Fig 3B) depending on the location of the spurious MUMs in other sequences.

**Figure 3:**
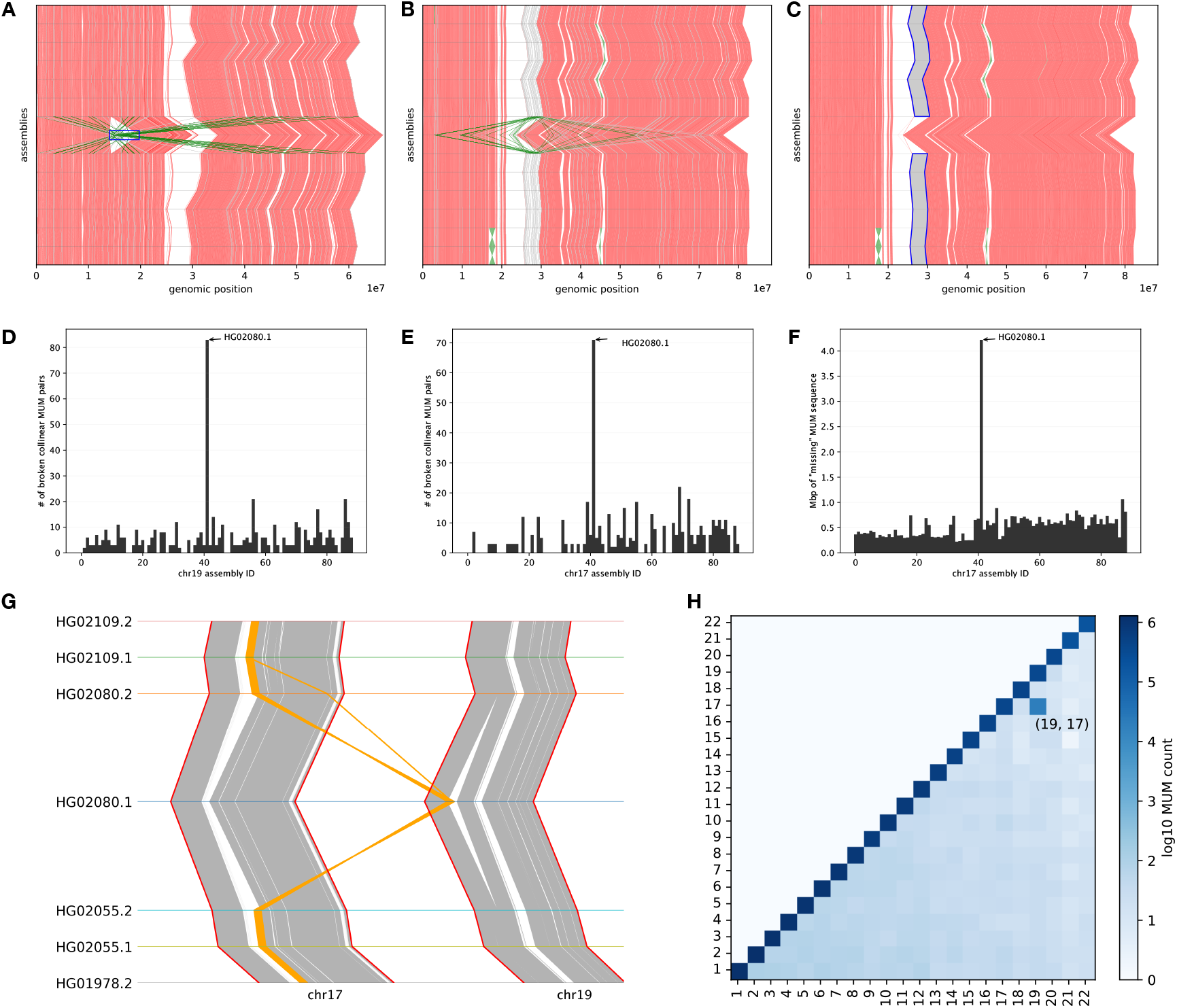
**(A-B)** MUM synteny visualization of collinear MUM blocks (red) and MUMs that break collinearity in a single sequence (gray (+)/green (-) based on orientation). **(C)** Large (>4 Mbp) syntenic region lost in HG02080.1, but recovered by partial MUMs (in gray). Evidence from non-collinear MUMs **(D-E)** and missing sequence present in partial MUMs **(F)** points a potential aberrant assembly artifact in the HG02080 paternal haplotype. **(G-H)** Genome-wide multi-MUMs reveal an interchromosomal join (confirmed to be a misassembly by HPRC [1]) in the aberrant regions.

We also found that Mumemto-reported partial MUMs (MUMs present in a subset of sequences) provide additional evidence. For instance, partial MUMs present in all but one sequence reveal large, private deletions (Figure 3C), which can indicate an assembly error or rare large-scale variant. Mumemto can identify these regions and quantify the sequence “missing” from each assembly using partial MUMs (Figure 3F).

As a case study, we used Mumemto to identify aberrant regions across HPRC haplotypes scaffolded using RagTag [21]. Figure 3 highlights a large insertion in the paternal haplotype of HG02080 chr19, identified by a spike in broken collinear MUM pairs. Additionally, we found a large deletion in chr17 of the same haplotype using partial MUMs. Computing multi-MUMs across full genome assemblies revealed a large interchromosomal join between chr17 and chr19 in the HG02080.1 assembly. This potential translocation was confirmed to be a misassembly by the HPRC team [1]. Identifying this problem was straightforward both quantitatively and visually using Mumemto multi-MUMs and partial multi-MUMs; it did not require finding pairwise alignments, building a graph, or computing a multiple alignment.

#### Scaffolding errors

We scaffolded the assembly contigs provided by HPRC using Rag-Tag, with default parameters and the T2T-CHM13 assembly as the reference [22]. Homology-based scaffolding is commonly used when there is no separate line of evidence such as Hi-C reads. However, scaffolding with respect to a single linear reference – even a high-quality reference – can bias contig placement and orientation [23], [24].

Mumemto can highlight potential scaffolding errors given the contig breakpoints in an assembly by identifying inversions at contig boundaries. We examined two instances of this on human chromosome 8 (Figure 5A). Both are located in one of the largest inversion polymorphisms in the human genome [25]. Based on RagTag scaffolding, the contigs covering the inversion in the paternal haplotypes of HG03098 and HG02148 are both oriented in the same direction as the CHM13 reference. RagTag assigns both contigs an orientation confidence score of 1.0, the highest possible confidence. However, it is clear from the MUM synteny visualization that both contigs should be reversed to preserve the flanking region orientation. This results in an inversion polymorphism that single reference-guided scaffolding would avoid. However, the overall pangenome synteny reveals that this polymorphism is common in the population. Mumemto synteny visualization and multi-MUM information can be used to correct the reference-guided scaffolding errors in each assembly using pangenome context.

### 2.4 Mumemto highlights pangenome-scale biology

#### Pangenomes across tree of life

We ran Mumemto on five recently released pangenome collections with genome lengths ranging from 13 Mbp (yeast) to 3 Gbp (human). For each, we computed all-pairs k-mer-based Jaccard similarities using Dashing2 [26]. As expected, the two interspecific datasets – potato [27], [28] and maize [29] – had the lowest inter-genome similarity, as captured by the inter-quartile range of Jaccard similarities (Table 2). The human pangenome had the highest inter-genome similarity, as well as the highest MUM coverage. We report the time and memory footprint for computing multi-MUMs across each pangenome dataset in Table 2. We also measured the wall-clock time required to compute MUMs over each set of chromosome assemblies. Since we used parallel threads for this, our memory measurement is the total memory footprint across all parallel threads.

**Table 2:** Multi-MUM and partial multi-MUM coverage, computation time, memory, and statistics for five pangenomes of varying sizes. `n/r` refers to the ratio of the total size of the pangenome (n) to the number of runs in the Burrow-Wheeler Transform over the sequence collection (r). The Jaccard index range is presented as an interquartile range (IQR), representing the 25th-75th percentile. Both Jaccard distance and `n/r` are included as approximate measures of intra-pangenome genomic divergence.

| dataset | N | # chr | genome size (Gbp) | $n/r$ | Jaccard index (IQR) | median MUM length (bp) | MUM coverage | median pMUM length (bp) | pMUM coverage | time (hh:mm:ss) | memory (GB) | ref |
| --- | --- | --- | --- | --- | --- | --- | --- | --- | --- | --- | --- | --- |
| Human | 89 | 22 | 2.86 | 135.16 | 0.92 – 0.93 | 110 | 83.76% | 445 | 91.70% | 4:14:11 | 707.5 | [1] |
| Maize | 27 | 10 | 2.13 | 42.33 | 0.44 – 0.47 | 41 | 13.48% | 139 | 30.36% | 2:29:49 | 661.6 | [29] |
| Potato | 60 | 12 | 0.752 | 30.20 | 0.32 – 0.45 | 29 | 10.31% | 51 | 62.54% | 1:38:50 | 545.5 | [27] |
| Arabidopsis | 69 | 5 | 0.135 | 42.03 | 0.56 – 0.63 | 37 | 43.06% | 91 | 73.88% | 32:53 | 92.6 | [30] |
| Yeast | 127 | 16 | 0.013 | 65.30 | 0.62 – 0.72 | 29 | 13.19% | 195 | 38.78% | 1:52 | 13.3 | [31] |

Table 2 also catalogs the strict (i.e. appearing in all sequences) MUM coverage and partial (i.e. appearing in a majority of sequences) MUM coverage. MUM coverage refers to the fraction of bases in each genome covered by at least one multi-MUM or partial MUM, averaged across all assemblies in the pangenome. Two datasets in particular, potato and *Arabidopsis*, displayed a large increase in coverage when including partial multi-MUMs. This trend generally indicates a small subset of sequences which form a distinct subgroup due to large genomic variation or incomplete assemblies within the group.

Mumemto can compute an outlier score when computing partial multi-MUMs. For a given sequence, this value is the aggregate of the lengths of partial MUMs in which the sequence is excluded, i.e. a high-scoring sequence tends not to share MUMs that are present in all other sequences. Figure 4 shows the outlier score for assemblies in the *Arabidopsis* (**A**) and potato (**B**) pangenomes. For potato, assemblies of *S. candolleanum*, a progenitor of cultivated potatoes which is considered a distinct clade within the Petota section of Solanum [27] tend to score higher, i.e. tend to be excluded from MUMs shared by others. Similarly for *Arabidopsis*, accessions from the African continent and from the geographically isolated Madeira islands have higher scores, along with two Asian accessions from Japan (Figure 4B).

**Figure 4:**
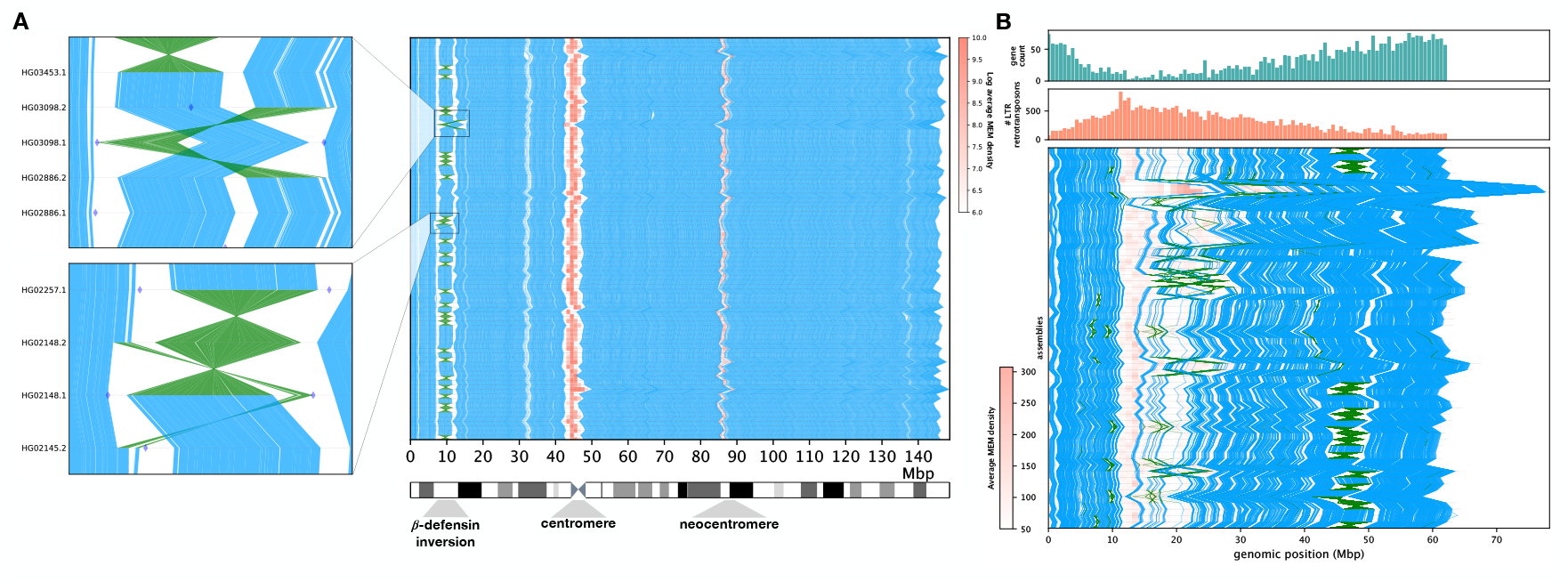
**(A)** HPRC chr8 assemblies visualized with multi-MUM synteny. Regions of high multi-MEM density shown in red. (Zoom panels) Two examples of incorrectly oriented contigs during scaffolding, with contig breakpoints represented by diamond markers. Inversions shown in green. **(B)** Assemblies of chr3 across the potato family, shown with multi-MUM synteny and MEM density colored in red. (top) Density of gene and LTR retrotransposon annotations for potato accession A6-26 (shown in the top row of syntenic view).

#### MUMs reveal genomic organization

Mumemto-computed multi-MUMs for potato chromosome 3 revealed a immediately noticeably pattern (Figure 5B). By visualizing the collinear blocks of multi-MUMs, we observed a denser arrangement of syntenic blocks conserved across the pangenome in the flanks of the chromosome. This correlates highly with gene density (Figure S2), as gene-rich regions are more evolutionarily conserved [32]. Regions with low MUM density also tend to be repetitive regions with more structural and less functional characteristics. The density of multi-MEMs (shown as a heatmap in red in Figure 5B) also recapitulates this trend, where spikes in multi-MEMs correspond to an increased density of LTR retrotransposons, the most abundant type of transposable element (TE) in the potato genome [32]. Both multi-MEMs and LTR TE density tends to be highest in the periocentromeric and centromeric regions, which has been previously observed in plants [32]. We note that this trend was clear despite the relatively low level of MUM coverage overall (Table 2).

**Figure 5:**
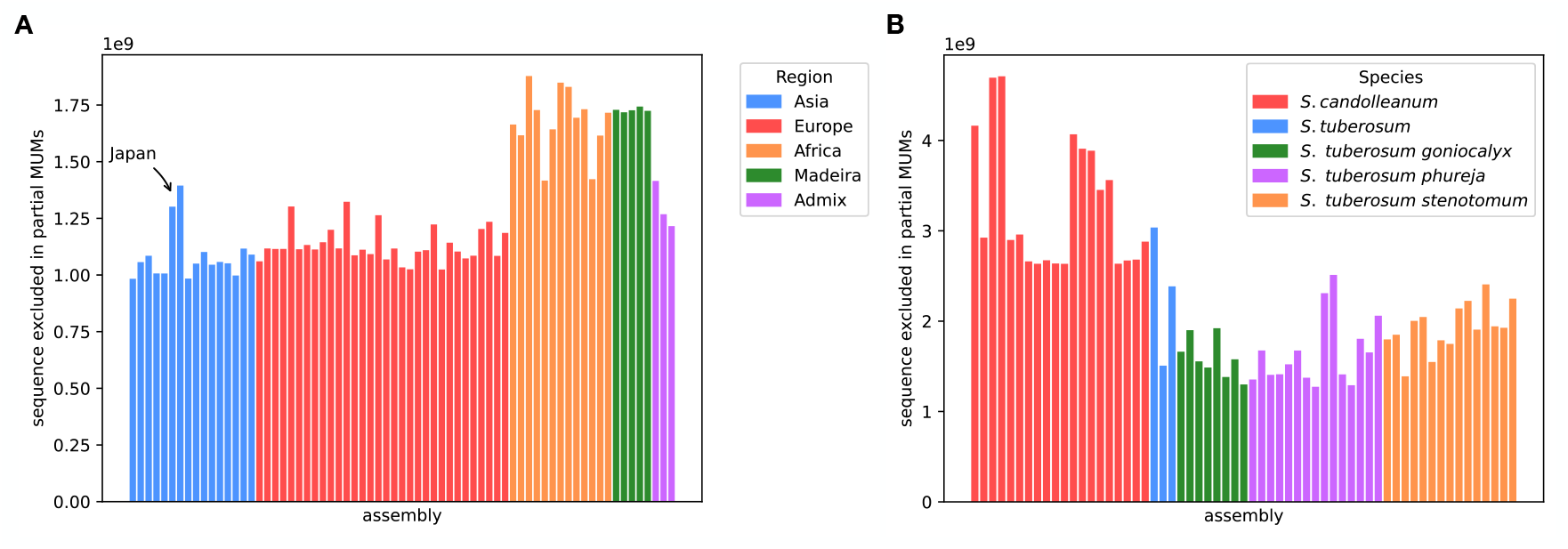
Aggregate length of partial MUMs that are not present in each genome assembly. **(A)** A. thaliana accesssions are grouped by geographical region, and **(B)** potato (Solanum section Petota) are grouped by species.

We also observed large-scale structural variations in chromosome 3. The largest of these was the ∼5.8 Mbp inversion polymorphism at a 40–50 Mbp offset in each assembly. Mumemto can further use the orientation of collinear multi-MUM blocks to identify large inversions, and can report approximate inversion boundaries. This inversion has been linked with the γ locus that controls tuber flesh color in potatoes and has been observed to cause suppressed recombination [27].

## 3 Discussion

Mumemto is an efficient tool for finding maximal exact matches, including multi-MUMs and related match types like partial MUMs and MEMs, across large collections of sequences. By computing these matches, Mumemto can rapidly define a pangenome coordinate system, aid in visualizing pangenome conservation and major structural variants, and reveal potential assembly issues and outliers. Mumemto also serves as an efficient multi-MUM-finding engine that can accelerate existing tools for core genome alignment and pangenome graph construction.

Mumemto can find partial multi-MUMs appearing in any subset of sequences as efficiently as it finds “strict” multi-MUMs. This enables a new view on pangenomes that reveals shared sequence within subgroups of a collection. Our findings for the *Arabidopsis* pangenome showed that partial MUMs can help to discern distinct subgroups of sequences and, ultimately, could quantify inter-sequence distances in a pangenome. Though this idea has been previously proposed [15], Mumemto’s algorithms allow for partial MUM computation at a scale of hundreds to thousands of genomes, potentially improving genomic distance and evolutionary inference for large sequence collections.

Mumemto computes multi-MUMs at the same time as PFP is computing the SA, LCP array, and the BWT. These arrays are exactly the key components of compressed full-text pangenome indexes like the r-index and move structure. These indexes can then be used to compute matching statistics [33] and other measures that quantify MEM-level similarity between query reads and pangenomes. In other work, we showed that considering multi-MUMs in the r-index improves read classification [34]. In this way, Mumemto can be integrated into a full-text indexing method to improve downstream alignment by providing a unifying coordinate system for full-text index-based alignment.

A potential drawback of Mumemto is its memory usage. Though PFP operates in compressed space, its memory footprint still reaches hundreds of gigabytes for large pangenomes. Currently, peak memory use could be reduced by splitting pangenomes by chromosome and computing only intra-chromosomal matches. Advances in compressed-space BWT construction [35] and improvements to prefix-free parsing [36] could help to decrease memory requirements without sacrificing inter-chromosomal resolution. We also plan to implement minimizer digestion to both decrease the input size as well as overcome minor genomic variation that may truncate syntenic blocks. Lastly, we plan to explore methods for merging multi-MUMs, such as the method in Parsnp2 [5] or index data structures like in RopeBWT3 [35], enabling a more incremental and parallelizable approach.

Due to how multi-MUMs are defined, they tend to cover less of the pangenome as the pangenome grows to include more individuals. While we find that multi-MUMs retain much of their utility even when they cover a lower percentage of the pangenome, we expect that a looser definition of MUM (e.g. a partial MUM present in a majority of sequences) becomes more appropriate as the pangenome grows. However, it will also be important to investigate other ways to loosen these requirements as the pangenome grows. Parsnp2, for example, uses a recursive process whereby it computes finer-grained multi-MUMs with respect to the spaces between previously identified coarser-grained multi-MUMs. We will explore a similar strategy in Mumemto, which could require multiple scans over the enhanced suffix array.

Visualization of pangenome synteny is currently limited to pairwise comparisons of genomes in a defined order. This order is arbitrary and thus multiple-sequence-based synteny is crucial to reveal the true pangenome coordinate system. Various methods exist for pairwise syntenic visualization [37], [38]. Although Mumemto implements a new visualization module intended for multi-MUMs, existing visualization methods could be used. However, this would require formatting the multi-MUM output as a set of pairwise comparisons for input, which would be computationally inefficient.

We showed that Mumemto can accelerate existing pipelines for pangenome alignment and construction. We also discussed the ability of Mumemto to reveal potential pangenome aberrations and misassemblies, improving newly assembled sequence collections and visualizing pangenomic variation structure. These use cases highlighted Mumemto’s potential as a core method for pangenomics, making it ideal as an initial tool in future pangenomic pipelines.

## 4 Methods

### Preliminaries

Given a text *T* of length *n, T* [*i*..*n*] is defined as the *i*th suffix. The suffix array (SA) over *T* is defined as the offsets of suffixes in *T*, ordered by lexicographic rank, such that *SA*[*i*] ≺ *SA*[*i* − 1]. The Burrows-Wheeler Transform *BWT*_*T*_ is a permutation of *T* defined such that *BWT*_*T*_ [*i*] = *T* [*SA*[*i*] −1], i.e. the BWT contains the characters preceding each suffix of *T* in lexicographic order. The Longest Common Prefix (LCP) array holds the lengths of the longest common prefixes between lexicographically successive suffixes. Formally, *LCP*_*T*_ [*i*] = *LCP* (*T* [*SA*[*i*]..*n*], *T* [*SA*[*i* − 1]..*n*]), where *LCP* [0] = 0. Here, we will consider that the SA, LCP array, and BWT are being constructed over a *collection* of sequences *T* = *T*_1_#*T*_2_# … *T*_N_, where # represents a unique sequence delimiter. We define the document array *DA*[*i*] as an array that identifies the sequence of origin (*T*_1_, *T*_2_, etc.) for the suffix at offset *i* in the SA.

#### Algorithm 1

Find multi-MEMs/MUMs

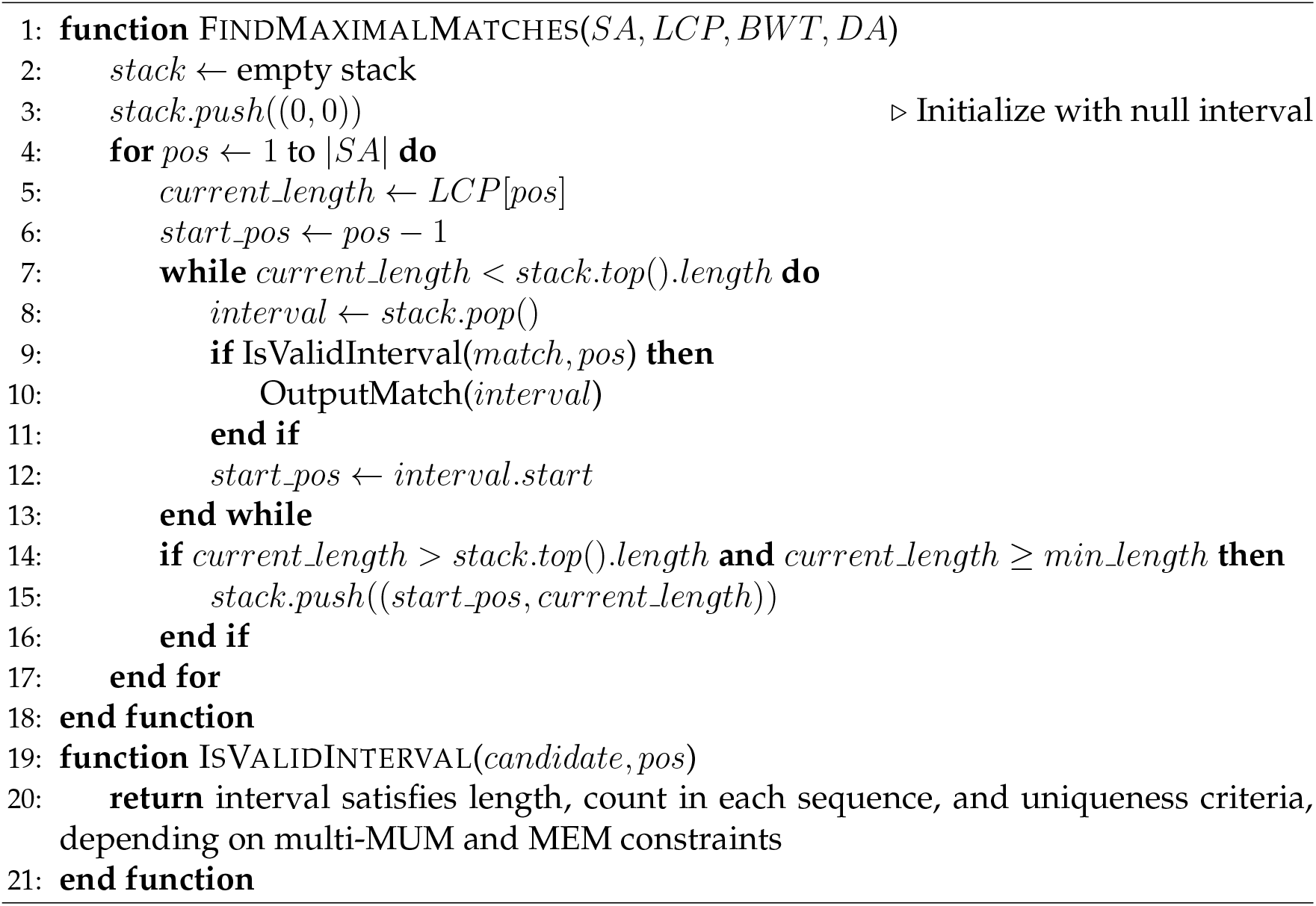

Multi-sequence maximal unique matches (**multi-MUMs**, referred to as simply MUMs when unambigious) are maximal exact matches that appear in all sequences (*T*_1_, *T*_2_, …, *T*_*N*_) and in each sequence exactly once. A maximal exact match is an exact match that cannot be extended further to the left or right. Multi-sequence maximal exact matches (**multi-MEMs**) relax the uniqueness constraint. That is, a multi-MEM appears in every sequence *at least* once, rather than *exactly* once. **Partial multi-MUMs** (**pMUMs**) are exact matches that appear exactly once in a subset of the sequences, and do not occur in the remaining sequences.

### Computing multi-MUMs and MEMs

Abouelhoda et al [39] showed how to use the SA and LCP array to simulate bottom-up traversal of a suffix tree and find supermaximal repeats between two sequences. Deogun et al [6] extended this to compute multi-MUMs by identifying LCP-intervals of size *N* that contain a suffix from each sequence and are not preceded by the same character. Note that the character that precedes the *i*^th^ suffix is *BWT*_*T*_ [*i*].

Mumemto’s core match-finding algorithm is adapted from bottom-up traversal ([3]; algorithm 4.1) and uses the multi-MUM properties defined by Deogun et al [6]. Its inputs are the SA, LCP array, BWT, and DA. We present this in Algorithm 1. By varying which properties we check in the `IsValidInterval` function, we can compute any form of multi-MUM or MEM using three main parameters: (1) number of sequences a match appears in (`-k`), (2) maximum occurrences within any given sequence (`-f`), (3) maximum total occurrences within the collection (`-F`) (see Figure 1, Figure S1). Algorithm 1 can be computed using just a buffer of values from each array, corresponding to the oldest interval in the stack, allowing for streaming computation. Extending this algorithm to compute matches on either strand of a nucleotide sequence, we include the forward and reverse complement of each sequence in the original text. A caveat is that the LCP-interval of a palindromic multi-MUM would appear as an interval of length 2*N*, violating the multi-MUM properties. Currently, Mumemto does not report palindromic multi-MUMs.

### Prefix-free Parsing

Typical suffix array construction algorithms (e.g. gSACA [40], [41]) scale linearly with input size and incur a memory footprint of many bytes per character of input sequence. Prefix-free parsing was introduced by Boucher et al. [9] as a method for computing a suffix array, BWT, and LCP array in compressed space, allowing it to scale to pangenomes, which tend to be highly repetitive. The key idea was to parse the input sequence into a dictionary *D* containing a set of phrases, and a parse *P*, holding the order in which the phrases must be concatenated to obtain the original text. Boucher et al. show that the BWT, LCP array, and SA can be computed in space proportional to *O*(|*D* + *P* |). For repetitive inputs, phrases will tend to be long (reducing |*P*|) and appear many times (reducing |*D* |). PFP computes the necessary arrays for Mumemto’s exact match computation. Importantly, it computes these values in order, allowing Mumemto to operate in a streaming fashion, and avoiding the need to store the SA and LCP fully in memory or on disk.

### Multi-MUM collinear blocks

We define an ordering of multi-MUMs in each sequence such that multi-MUM *C*[*i, j*] appears in sequence *i* with rank *j*. We define *collinear* multi-MUMs to be any multi-MUM that appears in a pair that is consecutive in every sequence, allowing for reversed pairs in case of a negative strand multi-MUM. A set of consecutive, collinear MUMs is defined as a collinear block. We refer to the region between collinear multi-MUMs as a collinear MUM gap. Collinear blocks can be optionally split into two when collinear MUM gaps are sufficiently large.

### Pangenome graph construction

Collinear multi-MUM blocks can be connected to form preliminary pangenome graphs. For each collinear block, the gaps between collinear multi-MUMs can be collapsed into the set of unique inter-MUM sequences. These represent a *snarl* [42] flanked by the multi-MUM sequence. Similarly, gaps between collinear blocks are collapsed when possible, and the necessary edges are included between MUM and inter-MUM nodes to provide haplotype walks that represent the input sequences. The nodes, edges, and walks are written to GFA format v1.1 for use with any downstream graph method.

## 5 Acknowledgments

We thank Todd Treangen and Bryce Kille for the valuable discussions on multi-MUMs. We also thank Katie Jenike for pointing us to useful pangenome datasets.

## 7 Availability of data and materials

Mumemto is available open source at https://github.com/vikshiv/mumemto, along with code for reproducing the results and figures.

## 8 Authors’ contributions

V.S.S. and B.L. conceived the project. V.S.S. wrote the software and conducted the experiments. V.S.S. and B.L. wrote the manuscript.

## 9 Ethics approval

Not applicable.

## 10 Consent for publication

Not applicable.

## 11 Additional Files

### 11.1 Supplement — Supplementary information

Contains Figure S1. PDF.

## References

[1] W.-W. Liao, M. Asri, J. Ebler, D. Doerr, M. Haukness, G. Hickey, S. Lu, J. K. Lucas, J. Monlong, H. J. Abel, et al., “A draft human pangenome reference,” Nature, vol. 617, no. 7960, pp. 312–324, 2023.

[2] G. Marçais, A. L. Delcher, A. M. Phillippy, R. Coston, S. L. Salzberg, and A. Zimin, “Mummer4: A fast and versatile genome alignment system,” PLOS Computational Biology, vol. 14, no. 1, e1005944, 2018.

[3] M. I. Abouelhoda, S. Kurtz, and E. Ohlebusch, “Replacing suffix trees with enhanced suffix arrays,” Journal of Discrete Algorithms, vol. 2, no. 1, pp. 53–86, 2004.

[4] A. C. Darling, B. Mau, F. R. Blattner, and N. T. Perna, “Mauve: Multiple alignment of conserved genomic sequence with rearrangements,” Genome Research, vol. 14, no. 7, pp. 1394–1403, 2004.

[5] B. Kille, M. G. Nute, V. Huang, E. Kim, A. M. Phillippy, and T. J. Treangen, “Parsnp 2.0: Scalable core-genome alignment for massive microbial datasets,” bioRxiv, pp. 2024–01, 2024.

[6] J. S. Deogun, J. Yang, and F. Ma, “Emagen: An efficient approach to multiple whole genome alignment,” in Proceedings of the second conference on Asia-Pacific bioinformatics, vol. 29, 2004, pp. 113–122.

[7] M. Höhl, S. Kurtz, and E. Ohlebusch, “Efficient multiple genome alignment,” Bioinformatics, vol. 18, no. suppl 1, S312–S320, 2002.

[8] T. J. Treangen and X. Messeguer, “M-gcat: Interactively and efficiently constructing large-scale multiple genome comparison frameworks in closely related species,” BMC Bioinformatics, vol. 7, pp. 1–15, 2006.

[9] C. Boucher, T. Gagie, A. Kuhnle, B. Langmead, G. Manzini, and T. Mun, “Prefix-free parsing for building big BWTs,” Algorithms for Molecular Biology, vol. 14, pp. 1–15, 2019.

[10] T. Gagie, G. Navarro, and N. Prezza, “Optimal-time text indexing in BWT-runs bounded space,” in Proceedings of the Twenty-Ninth Annual ACM-SIAM Symposium on Discrete Algorithms, SIAM, 2018, pp. 1459–1477.

[11] A. Kuhnle, T. Mun, C. Boucher, T. Gagie, B. Langmead, and G. Manzini, “Efficient construction of a complete index for pan-genomics read alignment,” Journal of Computational Biology, 2020.

[12] T. Nishimoto and Y. Tabei, “Optimal-time queries on BWT-runs compressed indexes,” in 48th International Colloquium on Automata, Languages, and Programming (ICALP 2021), Schloss-Dagstuhl-Leibniz Zentrum für Informatik, 2021.

[13] M. Zakeri, N. K. Brown, O. Y. Ahmed, T. Gagie, and B. Langmead, “Movi: A fast and cache-efficient full-text pangenome index,” iScience, vol. 27, no. 12, Dec. 2024.

[14] W. Beyer, A. M. Novak, G. Hickey, J. Chan, V. Tan, B. Paten, and D. R. Zerbino, “Sequence tube maps: Making graph genomes intuitive to commuters,” Bioinformatics, vol. 35, no. 24, pp. 5318–5320, 2019.

[15] A. E. Darling, B. Mau, and N. T. Perna, “Progressivemauve: Multiple genome alignment with gene gain, loss and rearrangement,” PLOS one, vol. 5, no. 6, e11147, 2010.

[16] H. Li, A collection of high-quality human assemblies, version v1.2, Zenodo, Oct. 2024. DOI: 10.5281/zenodo.13955431.

[17] T. J. Treangen, B. D. Ondov, S. Koren, and A. M. Phillippy, “The harvest suite for rapid core-genome alignment and visualization of thousands of intraspecific microbial genomes,” Genome Biology, vol. 15, pp. 1–15, 2014.

[18] G. Hickey, J. Monlong, J. Ebler, A. M. Novak, J. M. Eizenga, Y. Gao, T. Marschall, H. Li, and B. Paten, “Pangenome graph construction from genome alignments with Minigraph-Cactus,” Nature Biotechnology, pp. 1–11, 2023.

[19] J. Sirén, J. Monlong, X. Chang, A. M. Novak, J. M. Eizenga, C. Markello, J. A. Sibbesen, G. Hickey, P.-C. Chang, A. Carroll, et al., “Pangenomics enables genotyping of known structural variants in 5202 diverse genomes,” Science, vol. 374, no. 6574, abg8871, 2021.

[20] G. Baid, M. Nattestad, A. Kolesnikov, S. Goel, H. Yang, P.-C. Chang, and A. Carroll, “An extensive sequence dataset of gold-standard samples for benchmarking and development,” bioRxiv, pp. 2020–12, 2020.

[21] M. Alonge, L. Lebeigle, M. Kirsche, K. Jenike, S. Ou, S. Aganezov, X. Wang, Z. B. Lippman, M. C. Schatz, and S. Soyk, “Automated assembly scaffolding using Rag-Tag elevates a new tomato system for high-throughput genome editing,” Genome Biology, vol. 23, no. 1, p. 258, 2022.

[22] S. Nurk, S. Koren, A. Rhie, M. Rautiainen, A. V. Bzikadze, A. Mikheenko, M. R. Vollger, N. Altemose, L. Uralsky, A. Gershman, et al., “The complete sequence of a human genome,” Science, vol. 376, no. 6588, pp. 44–53, 2022.

[23] E. Bosi, B. Donati, M. Galardini, S. Brunetti, M.-F. Sagot, P. Lió, P. Crescenzi, R. Fani, and M. Fondi, “MeDuSa: A multi-draft based scaffolder,” Bioinformatics, vol. 31, no. 15, pp. 2443–2451, 2015.

[24] K.-T. Chen, C.-J. Chen, H.-T. Shen, C.-L. Liu, S.-H. Huang, and C. L. Lu, “Multi-CAR: A tool of contig scaffolding using multiple references,” BMC Bioinformatics, vol. 17, pp. 185–192, 2016.

[25] G. A. Logsdon, M. R. Vollger, P. Hsieh, Y. Mao, M. A. Liskovykh, S. Koren, S. Nurk, L. Mercuri, P. C. Dishuck, A. Rhie, et al., “The structure, function and evolution of a complete human chromosome 8,” Nature, vol. 593, no. 7857, pp. 101–107, 2021.

[26] D. N. Baker and B. Langmead, “Genomic sketching with multiplicities and locality-sensitive hashing using Dashing 2,” Genome Research, vol. 33, no. 7, pp. 1218–1227, 2023.

[27] D. Tang, Y. Jia, J. Zhang, H. Li, L. Cheng, P. Wang, Z. Bao, Z. Liu, S. Feng, X. Zhu, et al., “Genome evolution and diversity of wild and cultivated potatoes,” Nature, vol. 606, no. 7914, pp. 535–541, 2022.

[28] Potato Database, http://solomics.agis.org.cn/potato/, Accessed: 2024-12-10.

[29] M. B. Hufford, A. S. Seetharam, M. R. Woodhouse, K. M. Chougule, S. Ou, J. Liu, W. A. Ricci, T. Guo, A. Olson, Y. Qiu, et al., “De novo assembly, annotation, and comparative analysis of 26 diverse maize genomes,” Science, vol. 373, no. 6555, pp. 655–662, 2021.

[30] Q. Lian, B. Huettel, B. Walkemeier, B. Mayjonade, C. Lopez-Roques, L. Gil, F. Roux, K. Schneeberger, and R. Mercier, “A pan-genome of 69 Arabidopsis thaliana accessions reveals a conserved genome structure throughout the global species range,” Nature Genetics, pp. 1–10, 2024.

[31] S. O’donnell, J.-X. Yue, O. A. Saada, N. Agier, C. Caradec, T. Cokelaer, M. De Chiara, S. Delmas, F. Dutreux, T. Fournier, et al., “Telomere-to-telomere assemblies of 142 strains characterize the genome structural landscape in Saccharomyces cerevisiae,” Nature Genetics, vol. 55, no. 8, pp. 1390–1399, 2023.

[32] D. Zavallo, J. M. Crescente, M. Gantuz, M. Leone, L. S. Vanzetti, R. W. Masuelli, and S. Asurmendi, “Genomic re-assessment of the transposable element landscape of the potato genome,” Plant Cell Reports, vol. 39, pp. 1161–1174, 2020.

[33] O. Y. Ahmed, M. Rossi, T. Gagie, C. Boucher, and B. Langmead, “SPUMONI 2: Improved classification using a pangenome index of minimizer digests,” Genome Biology, vol. 24, no. 1, p. 122, 2023.

[34] N. K. Brown, V. S. Shivakumar, and B. Langmead, “Improved pangenomic classification accuracy with chain statistics,” bioRxiv, pp. 2024–10, 2024.

[35] H. Li, “BWT construction and search at the terabase scale,” Bioinformatics, btae717, 2024.

[36] E. Ferro, M. Oliva, T. Gagie, and C. Boucher, “Building a pangenome alignment index via recursive prefix-free parsing,” iScience, vol. 27, no. 10, 2024.

[37] D. Porubsky, X. Guitart, D. Yoo, P. C. Dishuck, W. T. Harvey, and E. E. Eichler, “Svbyeye: A visual tool to characterize structural variation among whole-genome assemblies,” bioRxiv, 2024.

[38] M. Goel and K. Schneeberger, “Plotsr: Visualizing structural similarities and rearrangements between multiple genomes,” Bioinformatics, vol. 38, no. 10, pp. 2922–2926, 2022.

[39] M. I. Abouelhoda, S. Kurtz, and E. Ohlebusch, “The enhanced suffix array and its applications to genome analysis,” in Algorithms in Bioinformatics: Second International Workshop, WABI 2002 Rome, Italy, September 17–21, 2002 Proceedings 2, Springer, 2002, pp. 449–463.

[40] G. Nong, “Practical linear-time o (1)-workspace suffix sorting for constant alphabets,” ACM Transactions on Information Systems (TOIS), vol. 31, no. 3, pp. 1–15, 2013.

[41] F. A. Louza, S. Gog, and G. P. Telles, “Inducing enhanced suffix arrays for string collections,” Theoretical Computer Science, vol. 678, pp. 22–39, 2017.

[42] E. Garrison, J. Sirén, A. M. Novak, G. Hickey, J. M. Eizenga, E. T. Dawson, W. Jones, S. Garg, C. Markello, M. F. Lin, et al., “Variation graph toolkit improves read mapping by representing genetic variation in the reference,” Nature Biotechnology, vol. 36, no. 9, pp. 875–879, 2018.

