## supplemental table and figures for "Mumemto: efficient maximal matching across pangenomes"

December 23, 2024

#### Supplementary Figures

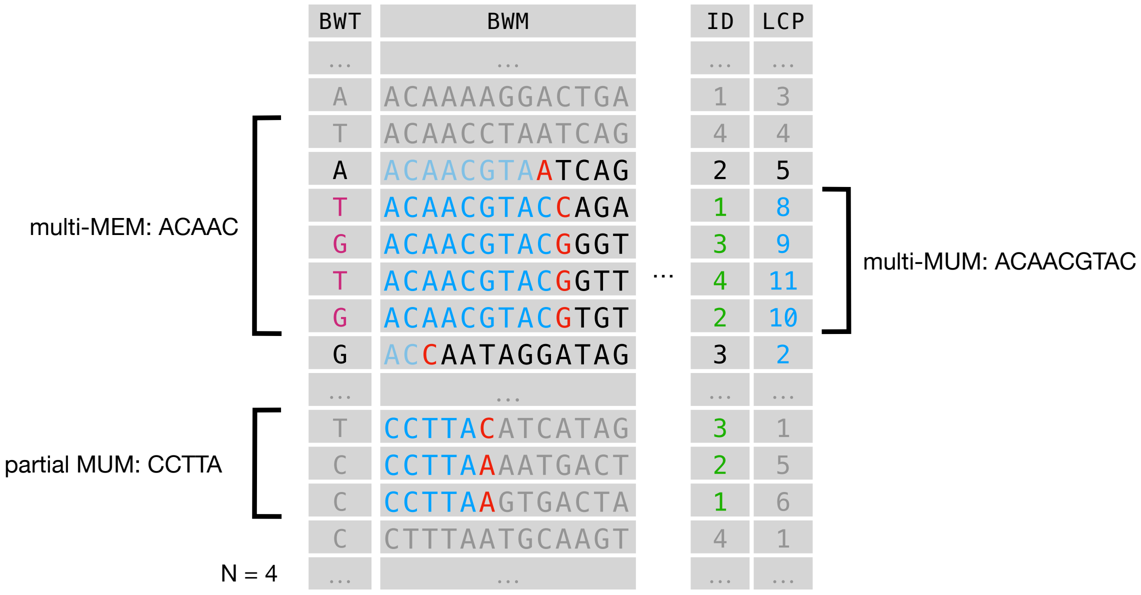


Figure S1: Example Burrows-Wheeler Matrix and transform (BWM / BWT), document array marking each suffix with the document of origin (ID), and longest common prefix array (LCP) for a set of four sequences. An example multi-MUM is highlighted, appearing in all sequences, along with a partial multi-MUM (appearing in only three sequences), and a multi-MEM (duplicated in sequences 2 and 4).

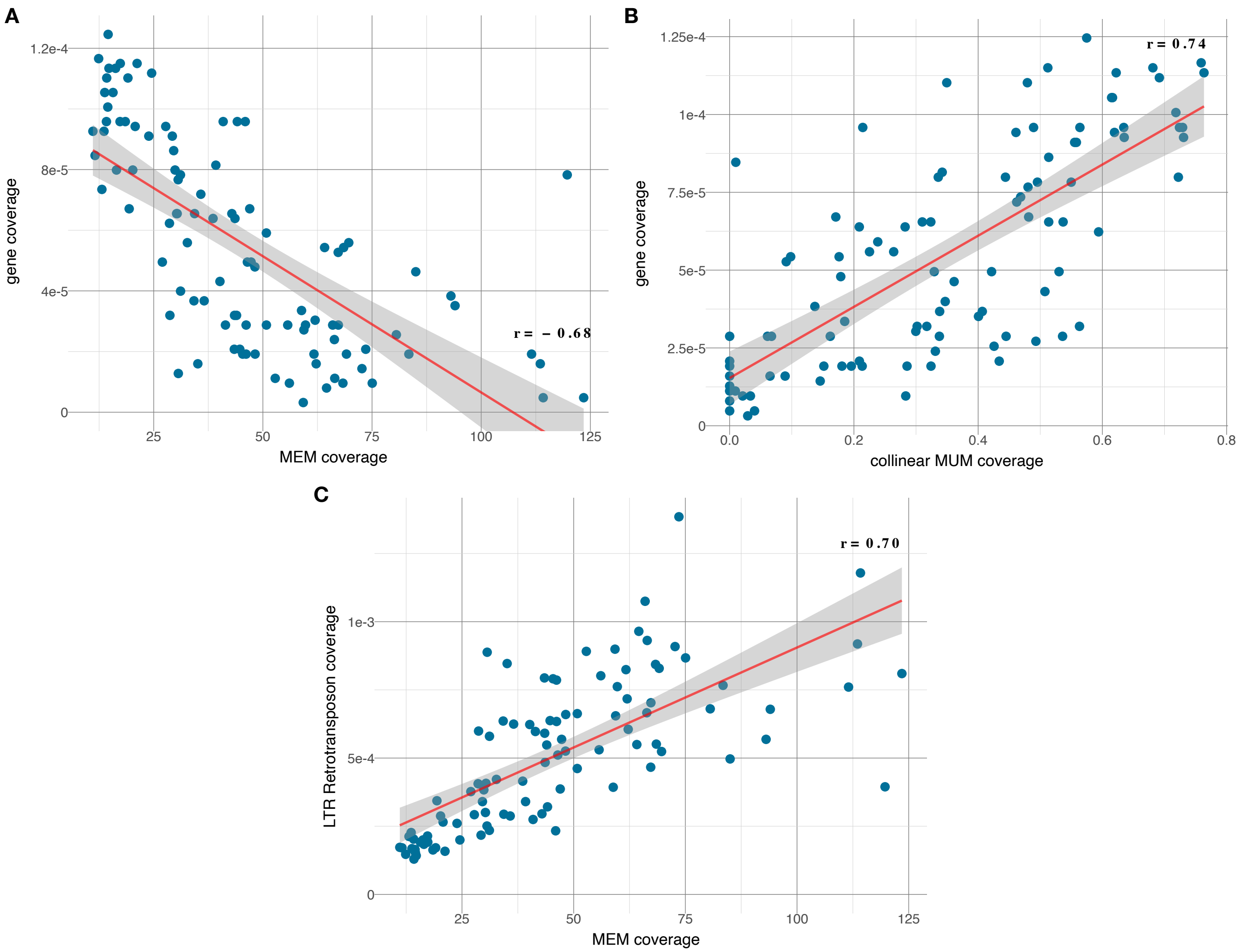


Figure S2: Correlation between **(A)** multi-MEM coverage and gene coverage, **(B)** collinear multi-MUM coverage and gene coverage, and **(C)** multi-MEM coverage and LTR retrotransposon coverage in local windows along the A6-26 assembly. Linear least-squares regression line and correlation reported for each comparison.

### Supplementary Tables

Table S1: Comparison of short read alignment with Giraffe against different pangenome graph indexes.

| | Minigraph-Cactus (MC) | Mumemto-full | Mumemto-collapsed | Mumemto + MC | single linear reference (CHM13) | diploid personalized reference (HG002) |
|---|---|---|---|---|---|---|
| aligned reads | 96922 | 96497 | 96726 | 97218 | 97316 | 97348 |
| perfect alignment | 75117 | 74496 | 74656 | 75495 | 65771 | 76081 |
| mean mapq | 54.1025 | 50.9781 | 51.6205 | 54.1671 | 55.8237 | 55.846 |
| speed (reads/s) | 1430.48 | 2343.37 | 3120.09 | 42.8307 | 6379.04 | 7627.51 |
| **Index sizes (MB)** | | | | | | |
| *.gbz | 114 | 339 | 225 | 153 | 44.8 | 49.8 |
| *.dist | 180 | 254 | 204 | 1271 | 27.2 | 34.7 |
| *.min | 713 | 1992 | 1360 | 759 | 601 | 605 |
